# TESTING THE USE OF XRCC6 GENE IN PHYLOGENETIC ANALYSES

**DOI:** 10.1101/2024.07.16.603754

**Authors:** Recep Çelik

**Affiliations:** Biology, Gazi University, ANKARA, 06560, Turkey

**Keywords:** Phylogenetic tree, XRCC6 gene, mtDNA, UPGMA, Neighbour-Joining

## Abstract

In this study, we investigated the potential use of the XRCC6 gene in the construction of phylogenetic trees used to visualise evolutionary relationships among species. XRCC6 gene is an important evolutionarily conserved gene that plays a role in DNA repair processes [1, 2]. The phylogenetic tree constructed with the XRCC6 gene using the Neighbour-Joining (NJ) algorithm was compared with the phylogenetic tree previously constructed with the UPGMA algorithm using mtDNA sequences. As a result of the comparison, it was observed that the trees constructed by both algorithms overlapped to a great extent in terms of topological similarity, support values of branches and species position. These findings indicate that XRCC6 gene can be used as a reliable genetic marker in phylogenetic analyses. The use of XRCC6 gene in phylogenetic analyses may contribute to a more comprehensive and accurate understanding of evolutionary relationships.

## 1. Introduction

Phylogenetic trees are important tools used to visualise evolutionary relationships between species [3]. These trees are constructed using genetic data and analysed with various algorithms. In recent years, mitochondrial DNA (mtDNA) analyses have formed the basis of phylogenetic studies[4]. However, the discovery of new gene candidates and the use of these genes in phylogenetic analyses can contribute to a more comprehensive understanding of evolutionary relationships. In this study, the use of XRCC6 gene as a potential candidate in phylogenetic tree drawing was examined. XRCC6 gene is an important gene involved in DNA repair processes and attracts attention with its evolutionarily conserved features [1, 2]. In our study, the phylogenetic tree constructed with the Neighbor-Joining (NJ) algorithm using the XRCC6 gene was compared with the UPGMA tree previously constructed with mtDNA analyses. This comparison aims to evaluate the usability of the XRCC6 gene in phylogenetic analyses.

## 2. Material and Method

Collection of Genomic Data:

The XRCC6 gene used in this study was obtained from the NCBI database. The gene sequences were selected for comparative analyses between various species. All sequences in the database were subjected to quality control and the appropriate sequences were used for phylogenetic analysis.

### Phylogenetic Tree Construction

Two different algorithms were used for phylogenetic analysis:

1. UPGMA (Unweighted Pair Group Method with Arithmetic Mean)[5]: This algorithm represents the phylogenetic tree previously constructed using mtDNA sequences. UPGMA visualises the relationships between species by calculating evolutionary distances[6].
2. Neighbour-Joining (NJ)[7]: This tree, constructed using the XRCC6 gene, determines relationships between species in a way that minimises evolutionary distances[8]. The NJ algorithm is known as a particularly fast and efficient method[9].

Phylogenetic trees for both algorithms were constructed using MEGA11 [10] software. Bootstrap analyses were performed to evaluate the accuracy of the trees and the reliability of each branch was indicated by percentage values.

### Comparison and Evaluation

The NJ tree and the UPGMA tree, which was previously generated by mtDNA analyses, were compared visually and statistically. The criteria used for comparison are as follows:

- Topological similarity: Similarity of the general structure of trees[11].
- Support values of branches : Similarity of branches with high support values in both trees.
- Position of species : Whether the species are in the same groups in both trees.

## 3. Findings

The first phylogenetic tree (Figure 1) was constructed with the UPGMA algorithm and the General Time Reversible (GTR) model using mtDNA sequences. In this tree, the evolutionary relationships between primates such as Homo sapiens, Pan troglodytes and Gorilla gorilla are clearly seen. The UPGMA tree shows close evolutionary relationships between species, with branches having high support values. In the UPGMA tree, the group “apes” (humans, chimpanzees, gorillas, orangutans[12]) is clearly separated by high support values. Likewise, the “OWM” (Old World monkeys[13]) and “NWM” (New World monkeys[14]) groups are also separated by high support values. The cladogram shows that mtDNA sequences accurately reflect evolutionary affinities.

**Figure 1.**
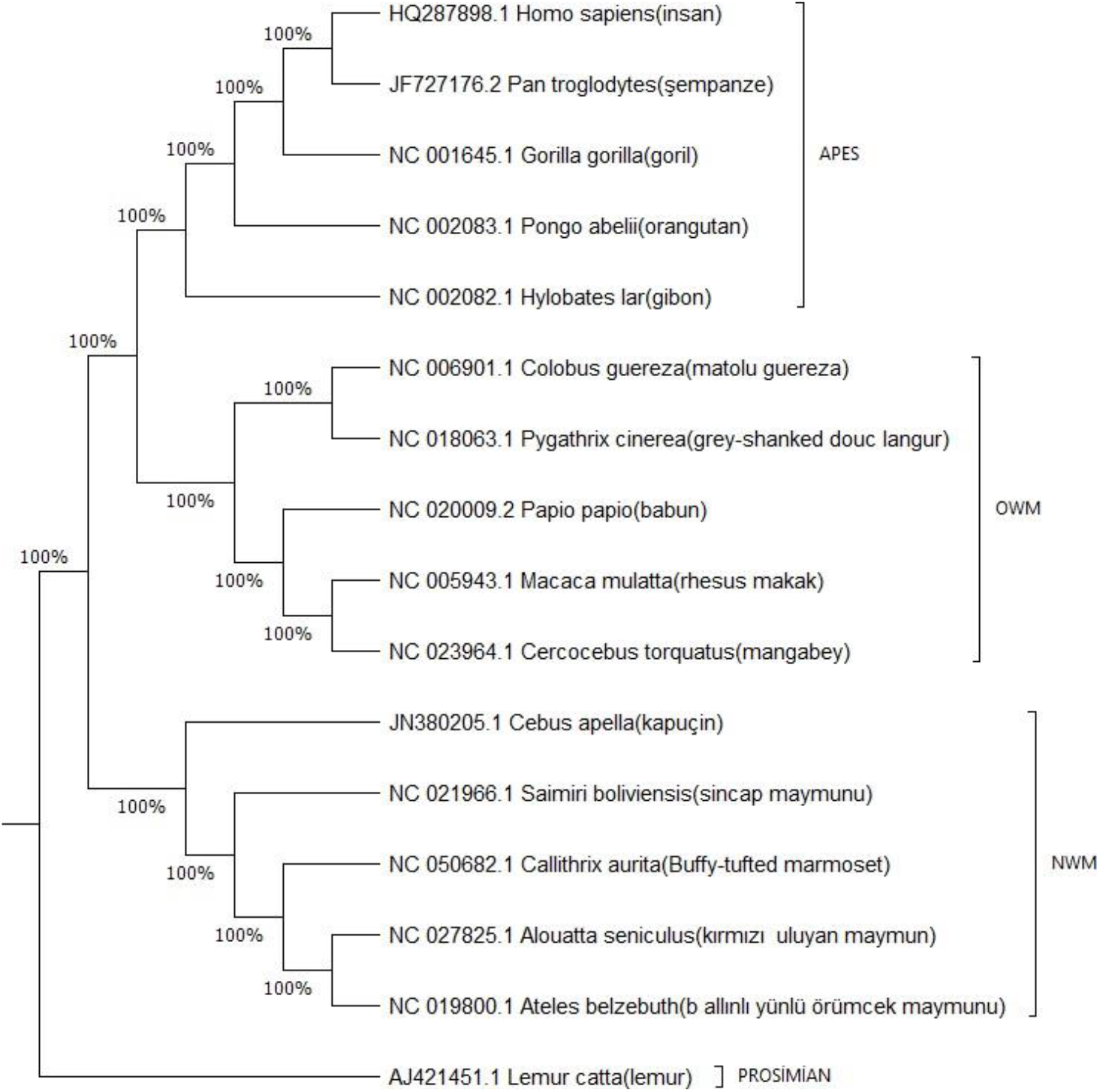
mtDNA sequences generated in MEGA11[10] using the UPGMA algorithm using mtDNA sequences

The second phylogenetic tree (Figure 2) was constructed with the Neighbor-Joining-General Time Reversible (GTR)+G model algorithm using the XRCC6 gene. Similar evolutionary relationships are observed in this tree. Species such as Homo sapiens, Pan troglodytes and Gorilla gorilla show similarly close evolutionary relationships. In the Neighbour-Joining tree, the “apes” group is clearly separated. Likewise, the “OWM” and “NWM” groups are also separated by significant support values. This phylogram shows the usability of the XRCC6 gene in phylogenetic analyses.

**Figure 2.**
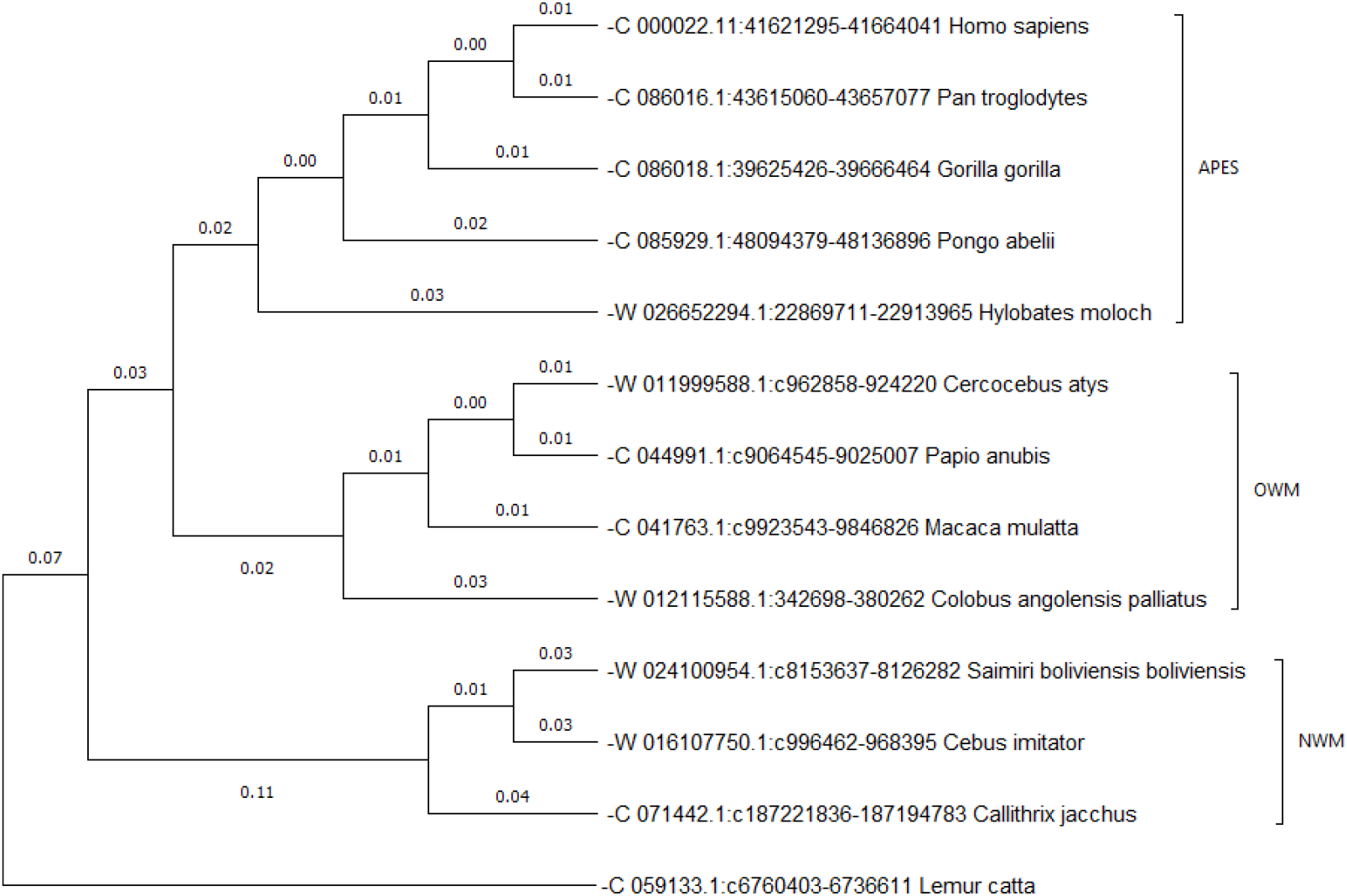
XRCC6 gene sequences were generated in MEGA11[10] using the Neighbor-Joining algorithm using sequences

## 4. Discussion and Conclusion

This study examined the utility of the XRCC6 gene in phylogenetic analyses and demonstrated that this gene is a reliable genetic marker. The Neighbor-Joining tree constructed using the XRCC6 gene gave similar results to the UPGMA tree constructed using mtDNA sequences. This indicates that the XRCC6 gene is a candidate gene region for phylogenetic analyses.

## Acknowledgements

I owe a debt of gratitude to “Regular Show” for motivating me all the time.

## Bibliography

[1] Jia, J., et al., Association Between the XRCC6 Polymorphisms and Cancer Risks: A Systematic Review and Meta-analysis. Medicine, 2015. 94(1).

[2] Thacker, J. and M.Z. Zdzienicka, The mammalian XRCC genes: their roles in DNA repair and genetic stability. DNA Repair, 2003. 2(6): p. 655–672.

[3] Yang, Z. and B. Rannala, Molecular phylogenetics: principles and practice. Nature Reviews Genetics, 2012. 13(5): p. 303–314.

[4] Sun, C.-H., et al., Mitochondrial Genome Structures and Phylogenetic Analyses of Two Tropical Characidae Fishes. Frontiers in Genetics, 2021. 12.

[5] Rohlf, F.J. and R.R. Sokal, Comments on Taxonomic Congruence. Systematic Zoology, 1980. 29(1): p. 97–101.

[6] Kalinowski, S.T., How well do evolutionary trees describe genetic relationships among populations? Heredity, 2009. 102(5): p. 506–513.

[7] Saitou, N. and M. Nei, The neighbour-joining method: a new method for reconstructing phylogenetic trees. Mol Biol Evol, 1987. 4(4): p. 406–25.

[8] Gascuel, O. and M. Steel, Neighbour-Joining Revealed. Molecular Biology and Evolution, 2006. 23(11): p. 1997–2000.

[9] Elias, I. and J. Lagergren, Fast neighbour joining. Theoretical Computer Science, 2009. 410(21): p. 1993-2000.

[10] Tamura, K., G. Stecher, and S. Kumar, MEGA11: Molecular Evolutionary Genetics Analysis Version 11. Molecular Biology and Evolution, 2021. 38(7): p. 3022–3027.

[11] Lopes, H. and M. Perretto, An Ant Colony system for large-scale phylogenetic tree reconstruction. Journal of Intelligent and Fuzzy Systems, 2007. 18: p. 575–583.

[12] Saber, M.M., et al., Emergence and Evolution of Hominidae-Specific Coding and Noncoding Genomic Sequences. Genome Biol Evol, 2016. 8(7): p. 2076–92.

[13] Elango, N., et al., Evolutionary rate variation in Old World monkeys. Biol Lett, 2009. 5(3): p. 405–8.

[14] Jacobs, G.H., New World Monkeys and Colour. International Journal of Primatology, 2007. 28(4): p. 729–759

